# Trivalent mRNA vaccine-candidate against seasonal flu with cross-specific humoral immune response

**DOI:** 10.1101/2023.12.30.573722

**Authors:** Elena P. Mazunina, Vladimir A. Gushchin, Denis A. Kleymenov, Andrey E. Sinyavin, Elena I. Burtseva, Maksim M. Shmarov, Evgenya A. Mukasheva, Evgeniia N. Bykonia, Sofia R. Kozlova, Elina A. Evgrafova, Anastasia N. Zolotar, Elena V Shidlovskaya, Elena S. Kirillova, Anastasiya S. Krepkaia, Evgeny V. Usachev, Nadezhda A. Kuznetsova, Igor A. Ivanov, Sergey E. Dmitriev, Roman A. Ivanov, Denis Y. Logunov, Alexander L. Gintsburg

## Abstract

Seasonal influenza remains a serious global health problem, leading to high mortality rates among the elderly and individuals with comorbidities. It also imposes a substantial economic burden through increased absenteeism during periods of active pathogen circulation. Vaccination is generally accepted as the most effective strategy for influenza prevention. As both influenza A and B viruses circulate and cause seasonal epidemics, vaccines need to include multiple antigens derived from different viral subtypes. While current influenza vaccines are effective, they still have limitations, including narrow specificity for certain serological variants, which may result in a mismatch between vaccine antigens and circulating strains. Additionally, the rapid variability of the virus poses challenges in providing extended protection beyond a single season. Therefore, mRNA technology is particularly promising for influenza prevention, as it enables the rapid development of multivalent vaccines and allows for quick updates of their antigenic composition. mRNA vaccines have already proven successful in preventing COVID-19 by eliciting rapid cellular and humoral immune responses. In this study, we present the development of a trivalent mRNA vaccine candidates, evaluate its immunogenicity using the hemagglutination inhibition assay, and assess its efficacy in animals. We demonstrate the higher immunogenicity of the mRNA vaccine candidates compared to the inactivated split influenza vaccine and its enhanced ability to generate a cross-specific humoral immune response. These findings highlight the potential mRNA technology in overcoming current limitations of influenza vaccines and hold promise for ensuring greater efficacy in preventing seasonal influenza outbreaks.

## INTRODUCTION

Seasonal influenza is a highly contagious respiratory disease caused by influenza A and B viruses that circulate around the world. Globally, influenza causes ∼1 billion cases of illness, 3 to 5 million cases of severe illness, and up to 500,000 deaths annually [1]. Pregnant women, children aged 6 months to 5 years, aged people (over 65 years of age), individuals with chronic disease, and healthcare workers are at increased risk of severe illness and serious complications from influenza virus infection [2,3]. A comprehensive study employing regression models revealed that the mortality rate associated with influenza between 1990 and 2017 was most pronounced among individuals over 70 years old, with a rate of 16.4 deaths per 100,000 (95% CI 11,6-21,9) [4]. Vaccination remains the primary strategy for reducing the incidence of influenza.

Various types of flu vaccines are available, including live attenuated, inactivated (whole virion, split, subunit), and recombinant vaccines [5]. The effectiveness of these vaccines (i.e. their ability to provide protection against influenza) may vary from season to season. At least two factors determine the likelihood of vaccine efficacy: (i) characteristics of the individual being vaccinated, such as age and health status, and (ii) the degree of matching between the vaccine composition and the influenza strains currently circulating in the human population [6]. Currently, most influenza vaccines are either quadrivalent (containing antigens of the H1N1 and H3N2 strains of influenza A combined with two lineages of influenza B, including the Victoria and Yamagata variants), or trivalent (containing the influenza A antigens of the H1N1 and H3N2 subtypes and one of the two influenza B subtypes) [7]. According to the CDC, in seasons when vaccine antigens matched circulating influenza viruses, vaccination reduced the risk of doctor visits related to influenza by 40% to 60% [8]. A 2021 study reported that among adults, vaccination was associated with a 26% lower risk of intensive care unit (ICU) admission and a 31% lower risk of death from influenza compared with those who were not vaccinated [9]. From 2010 to 2012, vaccination led to a 74% reduction in the risk of influenza-related ICU admissions in children [10], and according to a 2017 study, vaccination reduced the risk of influenza-related hospitalizations in older adults by an average of 40% between 2009 and 2016 [11]. Thus, vaccine prevention of influenza is both effective and justified.

In contrast, in instances where there was errors in selecting the appropriate antigenic composition, the efficacy of the vaccine was significantly compromised. An notable example was the 2017/18 vaccine, which exhibited low efficacy (∼25%) in the UK due to a mismatch with the predominant influenza A strain [11]. A similar decrease in effectiveness (to as low as 13%) relative to the H3N2 component of the vaccine was observed during the 2014-2015 season [12]. Throughout the history of influenza vaccination, there have been numerous occurrences of such mismatches, resulted in elevated rates of severe illness and mortality from influenza in certain seasons. To mitigate the impact of seasonal and pandemic influenza on public health, there is a need for vaccines that would offer broader and more reliable protection [13].

Different approaches and platforms have been employed in the development of new influenza vaccines, including virus-like particles, DNA/mRNA vaccines, baculovirus expression system, viral vectors, etc. Of particular interest are mRNA-based vaccines, which have demonstrated their efficacy and safety during the COVID-19 pandemic [14,15]. To date, a high immunogenicity of candidate mRNA influenza vaccines in animals and humans have been reported in a few studies [16–19]. In particular, immunization of mice with an mRNA candidate vaccine containing mRNAs encoding twenty hemagglutinins (HAs) of various influenza virus strains led to the formation of a prolonged humoral response to all twenty HAs [16]. Notably, the multivalent vaccine showed robust protection in animal models (mice, ferrets) when challenged with H1N1 influenza strains that varied in their similarity to the vaccine strain. The authors reported no mortality among vaccinated animals and observed a reduced disease severity (clinical scores) and a significantly lower weight loss compared to the control group [16].

A team led by G. Ciaramella has conducted extended preclinical and phase I clinical trials to evaluate the immunogenicity of modified mRNA encoding HA proteins from avian influenza viruses (H7 and H10) formulated in lipid nanoparticles (LNP) [19,20]. In mice, these vaccines demonstrated a 2-5-fold increase in hemagglutination inhibition assay (HAI) titers on day 21 after immunization, which remained at a consistent level throughout the year. The protective effect of the H7-mRNA was observed even with a minimal vaccine dose (0.4 μg per mouse), although it strongly depended on the period between infection and immunization (shorter intervals led to more rapid weight loss and their death from infection in vaccinated animals). Immunization of non-human primates with a single dose of 400 μg of H10- or H7-mRNA generated an immune response with HAI titers in serum ranging from 1:100 to 1:1 000; two weeks after repeated immunization, HAI titers reached 1:1 000 000. In two randomized, double-blind, placebo-controlled phase 1 studies involving healthy volunteers (N=201 for H10-mRNA and N=165 for H7-mRNA), the vaccines demonstrated favorable safety and reactogenicity profiles, as well as a robust humoral immune response [20]. Following double intramuscular immunization with 100 µg of H10-mRNA, all volunteers exhibited serum HAI titers exceeding 1:40, and 87% of participants showed microneutralization reaction titers of ≥1:20. For H7- mRNA, intramuscular administration of 10, 25, and 50 µg doses led to HAI titers exceeding ≥1:40 in ∼36%, 96%, and 90% of participants, respectively. At the same time, no significant HA-specific cellular immune response was observed in the IFN-γ ELISPOT assay [20].

Despite the growing body of research on the immunogenicity of mRNA vaccines against influenza, there is still a limited understanding of their potential to elicit a broad immune response against influenza strains with varying degrees of homology. In this study, we addressed this issue by presenting our own experience in the development of a trivalent mRNA flu vaccine and exploring its immunogenicity and protective efficacy in a mouse model. Through a two-dose immunization of mice, we observed not only a robust humoral immune response, but also cross-reactivity of this response against heterologous strains of the influenza virus.

## RESULTS

### Preparation and characterization of mRNA vaccine compositions

To develop the vaccine, HAs of three influenza viruses were chosen: A/Wisconsin/588/2019 (H1), A/Darwin/6/2021 (H3), B/Austria/1359417/2021 (IBV, Victoria lineage). These strains of influenza virus were included in the WHO recommendations for the 2022-2023 seasonal influenza vaccine in the northern hemisphere [21]. Сodon optimized DNA sequences of HA genes were synthesized and cloned into the pJAZZ-OK linear bacterial plasmid, as described previously [22]. *In vitro* synthesized mRNAs (schematically shown in **Figure 1**) included the cap-1 structure at the 5′ end; a 100-nt long poly(A)-tail at the 3′ end; the 5′ and 3′ untranslated regions (UTRs) from the human hemoglobin alpha subunit (*HBA1*) mRNA; and codon optimized coding sequences (CDS). N1-methylpseudouridines (m^1^) were co-transcriptionally incorporated into the mRNA instead of 100% uridines (U). mRNA-LNP formulations were prepared using the microfluidic NanoAssemblr Ignite mixer. The encapsulation efficiency was 89% (SD 1.2%) with a typical average particle size in the range of 68-71 nm with 0,105-0,148 polydispersity index (Supplementary Table S1).

**Figure 1.**
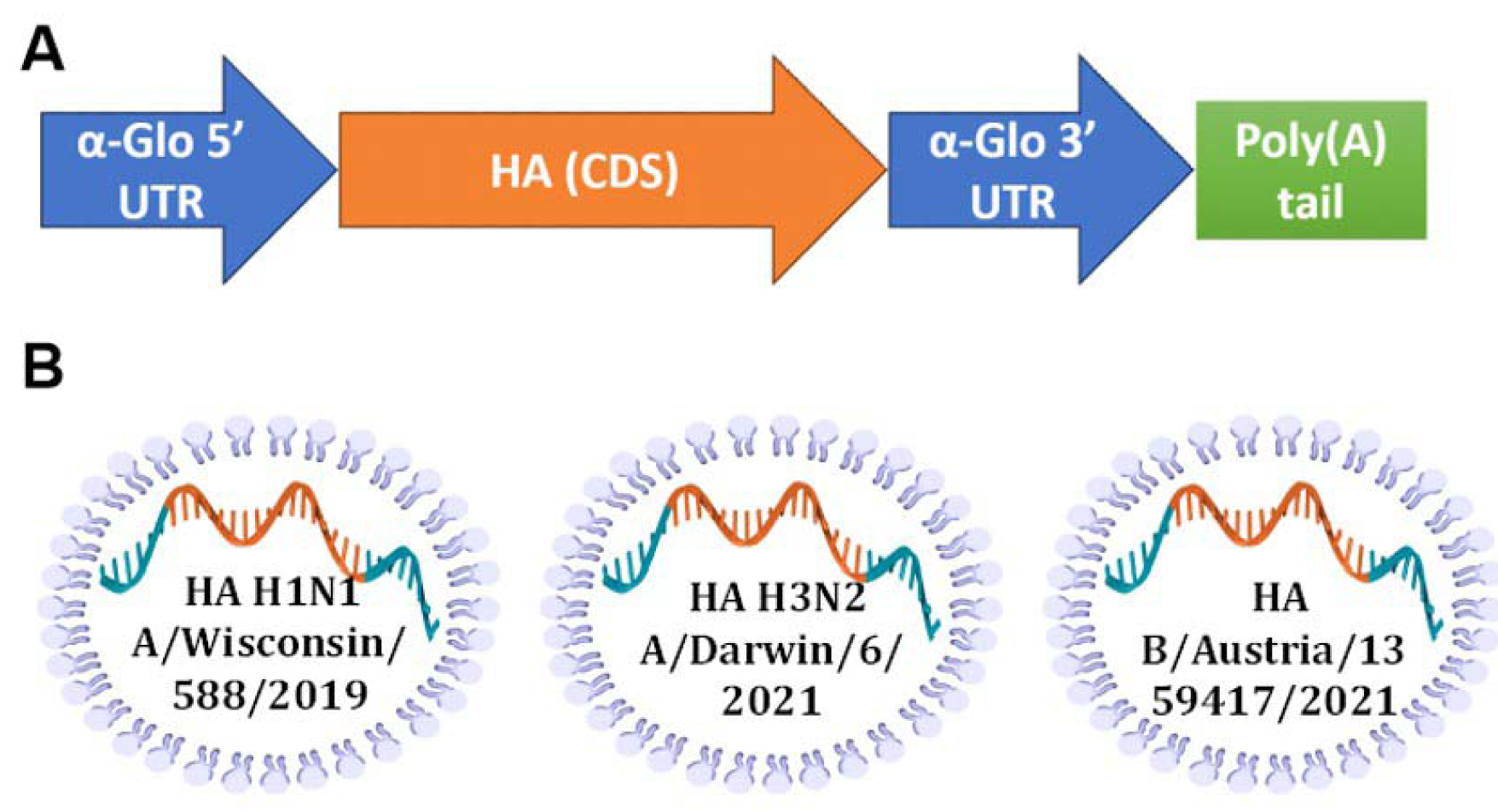
Design of mRNA-LNP for effective production of HA antigens in mammalian cells. A – scheme of mRNA with key parts, including 5’ and 3’ UTRs, influenza virus (IV) HA CDS, and poly(A)-tail; B – schematic visualization of separately formulated mRNAs encoding influenza HAs from three seasonal (2022-2023) vaccine strains.

A proper expression of the *in vitro* synthesized mRNAs was confirmed by transfection of cultured human embryonic kidney cells (HEK293) followed by their immunocytochemical staining for the H1N1 HA product. The presence of the HA protein (H1N1 A/Wisconsin/588/2019) was detected both inside the cells (intracellular staining, Figures 2A, C) and in the membrane-associated form (surface cell staining, Figures 2B, D). These results confirmed the translational activity of our synthetic mRNAs in cultured cells. In particular, the presence of the HA antigen on the cell surface indicated the correct design of the mRNA coding part containing a region for the HA transmembrane domain, which ensures the anchoring of the antigen to the host cell membrane.

**Figure 2.**
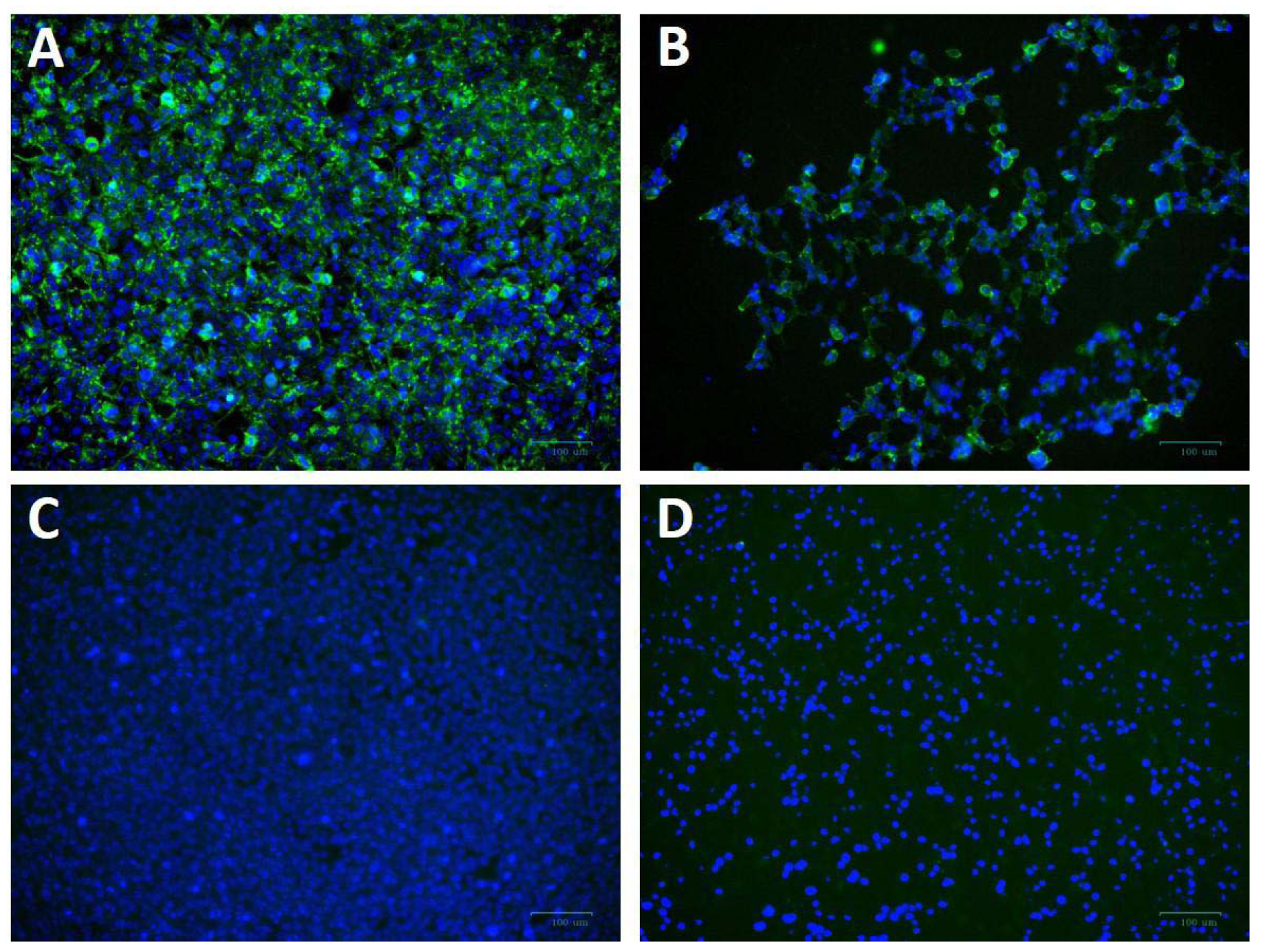
*In vitro* expression of hemagglutinin (H1N1 A/Victoria/2570/2019) in HEK293 cells transfected with synthetic mRNA. A, B – mRNA-transfected HEK293 cells were immunostained for intracellular (A) or surface HA protein. C, D – the same staining of untransfected HEK293 cells (C – intracellular, D – surface). Immunostaining was performed using primary polyclonal anti-Influenza A Antibody and Donkey Anti-Goat IgG H&L (Alexa Fluor® 488) secondary Ab (green), nuclei were stained with DAPI (blue). Scale bar 100 µM.

### Immunogenicity of two doses of the multivalent HA-mRNA vaccine

Immunogenicity of prepared mRNAs were investigated in mice using HAI assay, as described previously [23]. Initially, we tested the immunogenicity of H1 and H3 HA-encoding mRNAs separately (Supplementary Figure S1). HAI titers were measured after two 10-mg doses of individually formulated either H1 or H3 mRNA were administrated to BALB/c mice (n=6) intramuscularly (IM). HAI titers were up to 1:1280 above baseline by day 7 after second dose.

Next, to determine the immunogenicity of combined mRNA influenza vaccine candidate (mRNA-IV), HAI titers were determined in BALB/c mice (n=3) after two dose immunization with 15 µg of an equimolar mixture of individually formulated H1, H3, and IBV mRNA-LNPs. Thus, one dose of mRNA-IV contains 5 µg of each mRNA. This dose was selected as a potential 1/10 human dose of mRNA, by analogy with vaccines for the prevention of COVID-19. One control group of mice (n=3) received 6 µg (1/10 human dose) of split inactivated influenza vaccine (SIIV), another control group (n=3) received equal volume of sterile PBS. The SIIV contained 4 antigens according to WHO recommendations for the 2022-2023 seasonal influenza vaccine in the northern hemisphere [15]. The second dose was administered IM 14 days after the first dose.

#### H1 vaccine component

Analysis of the immune response to the H1 component of the mRNA vaccine showed high HAI titers against the homologous strain A/Wisconsin/588/2019 with geometric mean of 1:3413 (7 days after the second dose of vaccine). Compared to the control split vaccine (SIIV), the mRNA-LNP showed a rapid immune response significantly exceeding the limit of detection (LOD) as early as a week after the first vaccine dose. The average HAI titer against the A/Wisconsin/588/2019 antigen for the mRNA group after first dose was 1:126; for the SIIV group it was below the LOD (Mann-Whitney test *p=0.1*, Figure 3B). However, the Wilcoxon rank test showed no significant differences. On the 7th day after second vaccination, the differences in the level of antibody response in mRNA and split vaccine groups are especially noticeable - 1:3 225 and 1:40, respectively (80-fold differences), however, due to the small sample size, significant differences between these groups could not be found. Also, if we consider changes in the level of immune response within the group over time (on days 7, 21, and 35 after V1), a significant increase in the immune response is noted for the mRNA group after the second vaccine dose. For the mRNA group HAI titers against H1N1 on day 21 after V1 significantly exceed those on day 7 after V1 (*p=0.0429*, Friedman multiple comparison test). A decrease in antibody response by day 35 after V1 compared to day 21 was statistically not significant (Wilcoxon test *p=0.25*), the mean HAI titers differ by 2.2 times (Figure 3B). These differences are within the error of the HAI method, since the serum dilution factor in the study was equal to 2. A similar comparison in the SIIV group also did not reveal significant differences in HAI titers between 21 and 35 days against the A/Wisconsin/588/2019 strain (*p=0.75*). This may be due to the small sample size.

**Figure 3.**
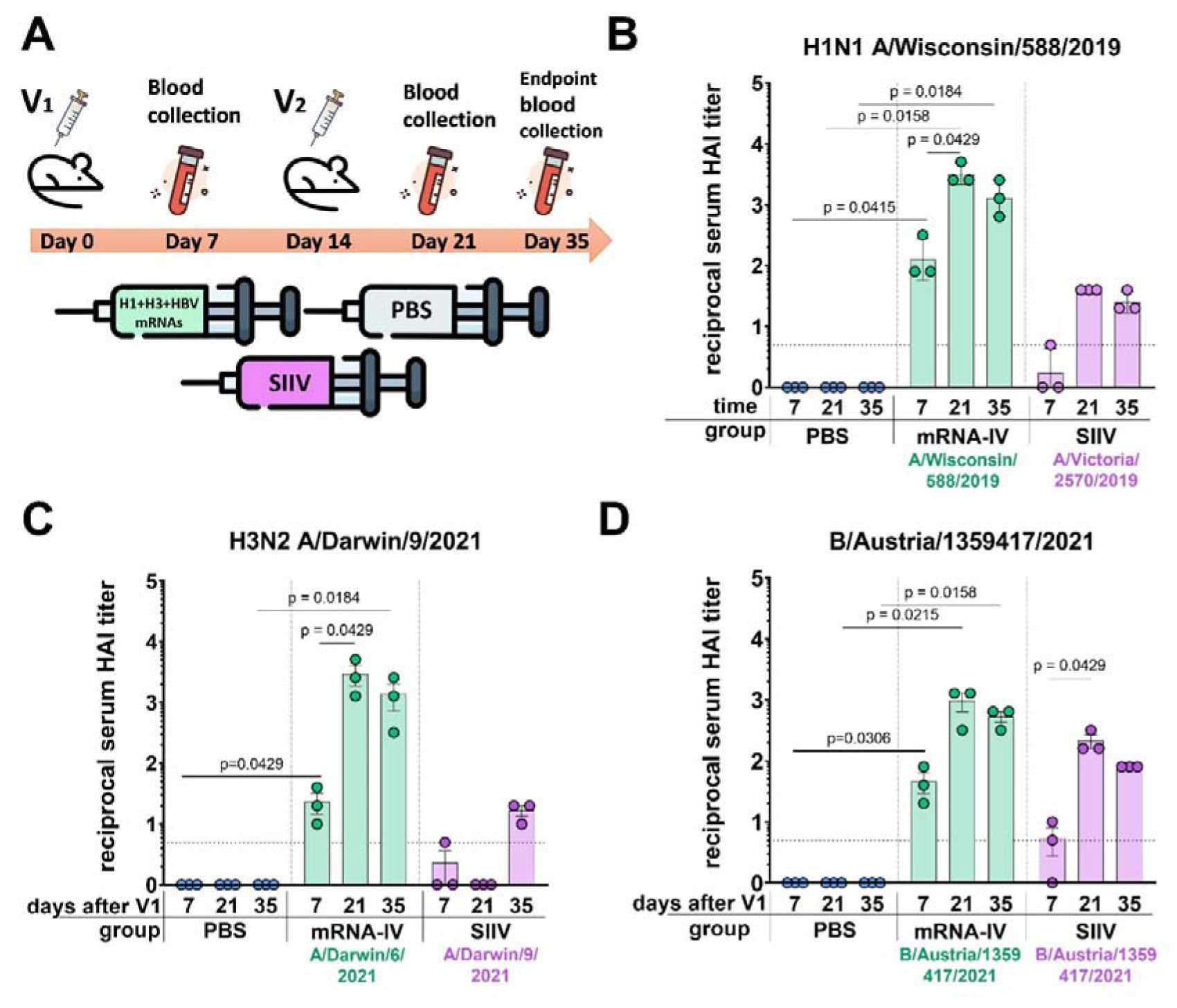
Mice immunized with mRNA-IV generate robust antibody responses. BALB/c mice (n=3 per group) were vaccinated intramuscularly at week 0 and 2 with a mixture of three different HA-mRNA-LNPs (a combined total dose of 15 µg mRNA per mouse (n=3), including 5 µg of each individual HA-mRNA-LNP). Control BALB/c mice were immunized intramuscularly with 1/10 human dose of quadrivalent split inactivated influenza vaccine (SIIV, a combined total dose of 6 µg per mouse, including 1.5 µg of each individual antigen), and another control BALB/c mice were administered PBS intramuscularly. A – scheme of the animal experiment. Serum HAI titers were determined after 7, 21, and 35 days post prime dose (V1) with antigens of vaccine strains: B – H1N1 A/Wisconsin/588/2019, С – H3N2 A/Darwin/9/2021, D – B/Austria/1359417/2021. Group mRNA-IV shown in green, SIIV – in violet, and PBS – in blue. Data are representative of one experiment and shown as geometric means ± SD. Data were analyzed using a Kruskal-Wallys test with Dunn’s multiple comparison test for inter-group analysis and Friedman test with Dunn’s comparison test for intra-group analysis of HAI titers through the time.

#### H3 vaccine component

For the H3 component of the vaccines, we obtained a similar immunological profile, but with more pronounced differences between mRNA and split vaccine in terms of the level of immune response (Figure 3C). As we did not have the influenza vaccine strain A/Darwin/9/2021 in our virus strain collection, HAI titers were determined using the antigen of serologically close strain - A/Darwin/6/2021 (2 amino acids substitutions in HA protein sequence - G69D, D202N). Thus, H3-mRNA (A/Darwin/9/2021), in trivalent mRNA-IV vaccine caused a detectable antibody response already on day 7 after V1 (mean 1:23), while for split vaccine this value was below the LOD (Figure 3C). Seven days after the second vaccination, the mean titer of HAI in the mRNA group reaches 1:2 987, while in sera from mice that received the split inactivated vaccine, this value remains below the LOD (Figure 3C). Increase of HAI titer in serum of mice vaccinated by SIIV was observed only to 35 days after V1, however it remains 86-fold lower than this value for the mRNA vaccine (mean HAI titer in mRNA vaccinated mice - 1:1 387). When comparing immunogenicity through the time of experiment (7, 21, and 35 days after V1) within the mRNA group, there was observed significant increase after the second dose of mRNA on day 21 (with a mean titer of 1:2 987), in comparison with serum titers on day 7 (*p=0.0429*, Friedman multiple comparison test). There were no significant differences in titers between days 21 and 35 (*p=0.66*).

#### B (Victoria) vaccine component

The immune response in vaccinated mice to influenza B virus (Victoria lineage) was lower compared to two previous vaccine components (H1 and H3). Two-dose vaccination with mRNA-LNP resulted in lower serum HAI titers to B/Austria/1359417/2021 antigens than to H1N1 antigens (mean HAI titer in mRNA vaccinated mice – 1:960 at 21 days post V1, Figure 3D). However, just 7 days after the first dose, HAI titers for the mRNA group exceeded the LOD (mean HAI titer - 1:45), while in the split vaccine group, the mean HAI titer was below the LOD (mean HAI titer - 1:5). The difference in the level of immune response to mRNA and split vaccines is much less for the B component of the vaccine than for the H1 and H3 components. A decrease in antibody response by day 35 after V1 compared to day 21 was not statistically significant (*p=0.25*, Wilcoxon test).

### Cross-reactivity of immune response 3 weeks after two-dose vaccination

On day 35 after the start of the experiment, the whole blood was collected from the mice for more in-depth studies. This allowed us to study the cross-reactivity of the immune response after the vaccination to more than one different influenza strain only for endpoint serum sample of mice (21 days post second dose, Figure 4). For the H1N1 strains we used (Wisconsin/588/2019, Victoria/2570/2021, Guangdong-Maonan/SWL1536/2019, Moscow/52/2022, California/07/2009), the immune response is highly cross-reactive in the group of mRNA-vaccinated animals. The geometric mean HAI titers against H1N1 strains were from 1:806 (against Guangdong-Maonan/SWL1536/2019 antigen) to 1:2 032 (Figure 4A). There was no significant reduction in HAI titers even to distant pandemic A/California/07/2009 strain with mean value 1:1 015 (Figure 4A). The immune response to SIIV was not as robust as to mRNA-IV, the mean titer value did not exceed 1:31 (against Victoria/2570/2021 antigen), which more than 60-fold lower comparing with mRNA. For single H1-mRNA study of cross-reactivity of immune response showed similar results (Supplementary Figure S2).

**Figure 4.**
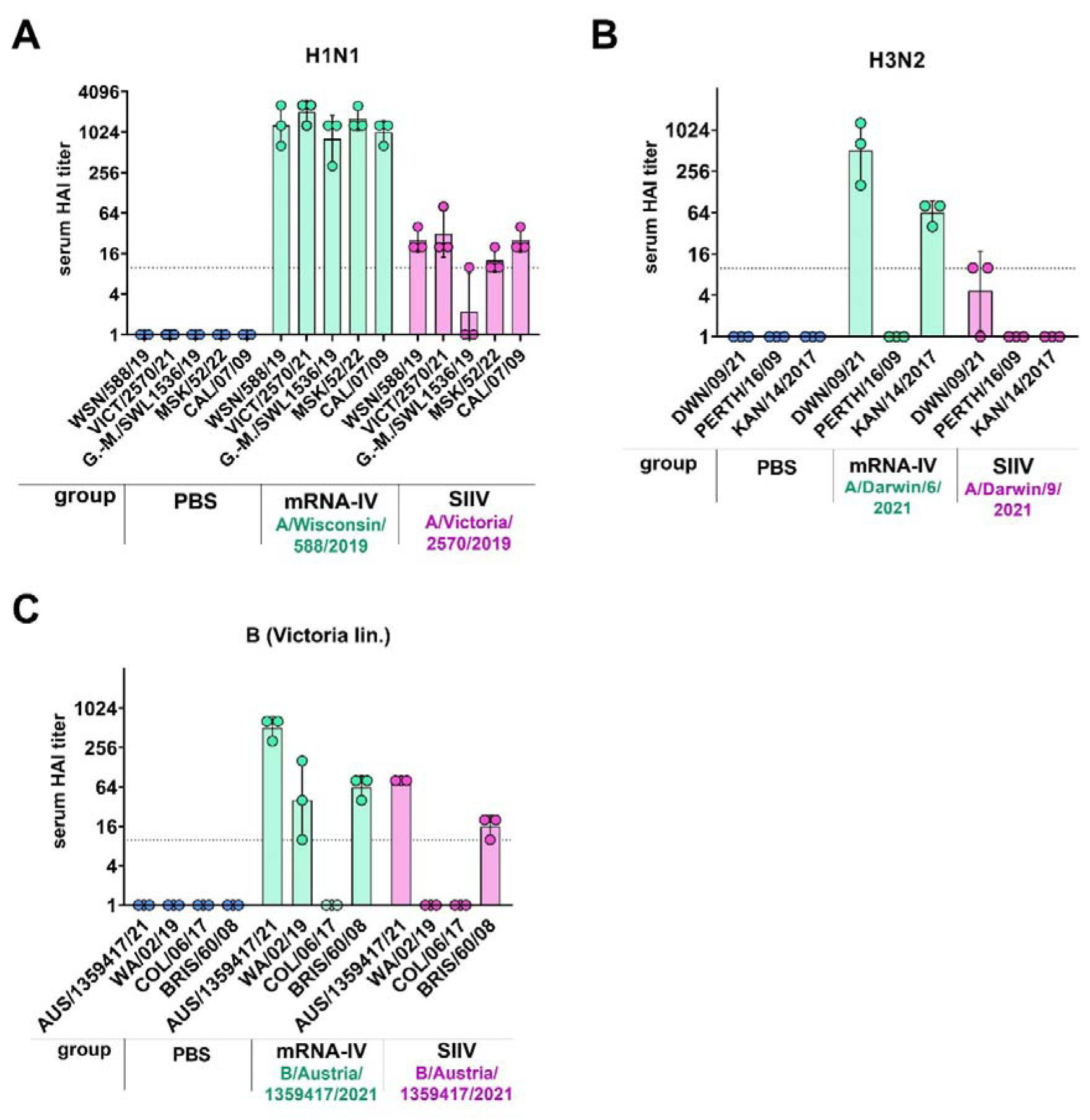
Mice immunized with mRNA-IV generate cross-reactive antibody responses. BALB/c mice (n=3 per group) were vaccinated intramuscularly at week 0 and 2 with 3 different HA mRNA-LNPs (a combined total dose of 15 µg per mouse (n=3), including 5 µg of each individual HA mRNA-LNP). Control BALB/c mice were immunized intramuscularly with 1/10 human dose of registered quadrivalent split inactivated influenza vaccine (SIIV, a combined total dose of 6 µg per mouse, containing 1.5 µg of each individual antigen), and another control BALB/c mice were administered PBS intramuscularly. Serum titers in heterologic HAI were determined after 35 days post prime dose with antigens of different influenza strains: A – strains of H1N1 influenza subtype (Wisconsin/588/2019, Guangdong-Maonan/SWL1536/2019, Moscow/52/2022, California/07/2009), B – strains of H3N2 influenza subtype (Darwin/9/2021, Perth/165/2009, Kansas/14/2017), С – strains of influenza B (Victoria lineage – Austria/1359417/2021, Washington/02/2019, Colorado/06/2017, Brisbane/60/2008). Group mRNA-IV shown in green, SIIV - in violet and PBS - in blue. Data are representative of one experiment and are expressed as geometric means ± SD.

The cross-reactivity of immune response to H3 component of the vaccine was manifested by decreased titers to the Kansas/14/2017 strain (the geometric mean HAI titer was 1:63) and no detectable response to the Perth/165/2009 strain, while response to serologically close strain A/Darwin/6/2021 remained to be high with geometric mean HAI titer 1:1 015.

Noticeable results were obtained in a heterologous HAI test with the antigens of influenza virus B strains (B/Washington/02/2019, Colorado/06/2017 and Brisbane/60/2008, Figure 3C). Immune response against listed strains was reduced compared to homologous HAI results (Figure 3C) for both mRNA and split inactivated vaccine. In the mRNA group, the decrease in mean HAI titers to the Washington/02/2019 antigen was 12-fold and to the Brisbane/60/2008 antigen was 8-fold. No immune response to the Colorado/06/2017 antigen was detected in the serum of mice vaccinated with both mRNA and SIIV. In the SIIV group, no immune response to the Washington/02/2019 antigen was detected and the reduction in HAI titers to Brisbane/60/2008 compared to homologous was more than 5-fold.

### Protective efficacy of two doses of the trivalent HA-mRNA vaccine

In order to evaluate the effectiveness of immunity formed by mRNA-LNP vaccination, we conducted an experiment using an animal model of lethal influenza infection. This model included infection of animals vaccinated according to the chosen regimen with mouse-adapted influenza A/Victoria/2570/2019 virus. The design of the experiment is shown in Figure 5A.

**Figure 5.**
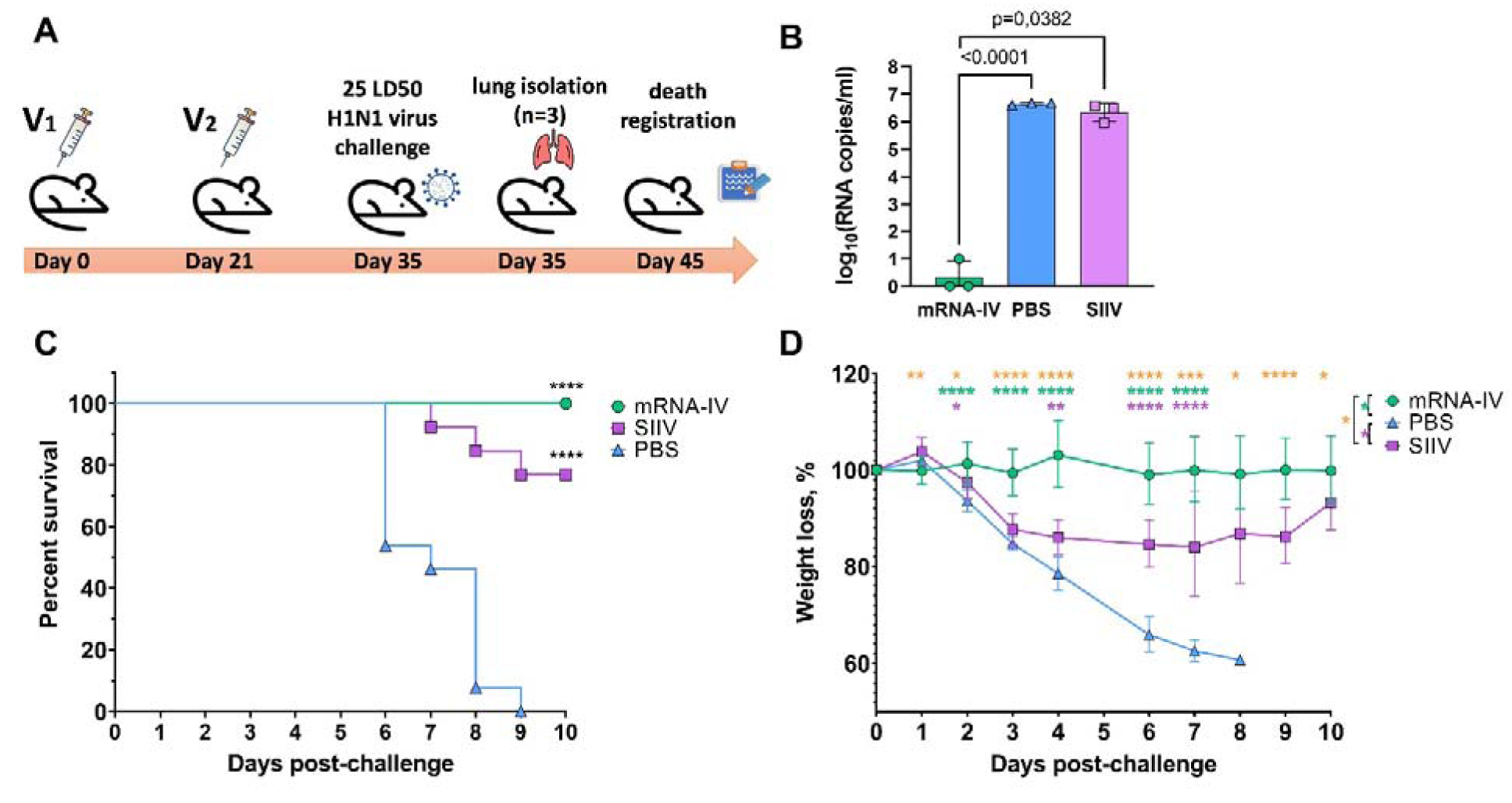
The mRNA-IV vaccine protects mice from influenza infection with antigenically close H1N1 strain. BALB/c mice were immunized with two-dose of mRNA vaccine and challenged with 25 LD_50_ of H1N1 virus (A/Victoria/2570/2019, mouse adapted). A - scheme of challenge experiment. At third day post infection lungs of mice (n=3) from every group were harvested and analyzed by qPCR for viral load determination (B). Survival (C) and weight loss (D) of the remaining animals were recorded post challenge. Data are representative of one experiment and shown as geometric means ± SD. Data were compared using a Long-rank (Mantel-Cox) test with Dunn’s multiple comparison test for survival analysis and Tukey’s multiple comparisons test for analysis of weight loss differences between groups. The viral load data were compared using Mann-Whitney test. P-value shown as asterisks, *\*p < 0.05, **p < 0.01, ***p < 0.001, ****p < 0.0001*.

Animals immunized with PBS (n=13) and animals that received 1/10 human dose of inactivated split vaccine were used as comparison groups. Fourteen days after second immunization according to the scheme in Figure 5A mice were infected intranasally with 25 LD50 of H1N1 pdm09 virus (A/Victoria/2570/2019, mouse adapted), which had 99,8% HA amino acid homology to the H1 component of the mRNA vaccine. On the third day after infection, 3 mice from each group were euthanized for lung isolation. Viral load was assessed in lung homogenates by RT-PCR with normalization to a housekeeping gene, *PDK1* (encoding pyruvate dehydrogenase kinase-1). In the mRNA vaccinated mice group, the average viral load in the lungs was 3.5 copies/ml, while in the PBS and SIIV control groups, the mean Log10 viral load was 6,65 and 6,4 genome copies/ml, respectively (Figure 5B). The viral load in the lungs of mRNA vaccinated mice was almost undetectable in contrast to the PBS and SIIV control groups (*p<0.0001* and *p=0.0382*, respectively, Figure 5B). On day 6 post-infection, 46% of the animals in the control group died and all animals died on day 9 (Figure 5C). In the group of animals that received the inactivated split vaccine, the lethality rate on (or between) days 7–9 of infection was 23%. No animal deaths were observed in the group that received the mRNA-IV. Both vaccines protected animals from 25 LD_50_ doses of virus with statistical reliability (*p<0,0001* compared to the PBS group). No significant differences were found between vaccines according to these statistical tests. Evaluation of weight dynamics shows that animals from the control group started active weight loss from the second day after infection (Figure 5D). In the group of animals receiving SIIV, weight loss was observed until the 3rd day of infection with subsequent recovery. No weight loss was observed in the group of animals receiving mRNA. On day 1 after infection, an average increase in body weight of 3% was observed for animals in the SIIV group and the value of this parameter was significantly higher than that of the mRNA-IV group. However, from day 2 post-infection until the end of observations (day 10), weight loss was minimal for the mRNA-IV group and was significantly lower for the other two groups of mice that received inactivated vaccine or PBS (Figure 5D).

## DISCUSSION

Due to the ongoing genetic perturbations and evolution of influenza viruses, including antigenic drift [24] and recombination [25,26], the antigenic characteristics of the pathogen undergo continuous changes [24]. As a result, the immunity generated by previous exposure or vaccination becomes less effective. This continually puts the human population at risk of seasonal influenza epidemics. To address this challenge, the WHO Global Influenza Surveillance and Response System (GISRS) monitors influenza viruses circulating in humans and updates the antigenic composition of vaccines twice a year (for the southern and northern hemispheres) [27]. Manufacturers of influenza vaccines receive guidance on the specific antigenic composition, including two serologic variants of influenza A and two serologic variants of influenza B, to be incorporated in vaccines for the upcoming season [28]. However, mistakes in the strain selection, as well as the need to enhance protection in vulnerable groups, have prompted the search for novel approaches to develop influenza vaccines that offer broader and more effective protection [29,30].

The mRNA platforms hold significant promise for the development of influenza vaccines and offer several advantages. First, it enables rapid vaccine creation and production in event of a completely new virus strain, allowing for a more timely response to emerging threats [31]. Second, unlike traditional methods, mRNA vaccine production does not require the use of live virus and can be based on synthetic sequences obtained in the laboratory. This eliminates the risk of unintended mutations that may arise from virus passages. Third, mRNA-based vaccines are highly precise in targeting the immune response as they express only specific antigens, such as influenza HA [32]. Finally, the delivery of mRNA formulated in LNPs not only promotes both humoral and cellular immune responses but also activates the innate immune system, further enhancing the effectiveness of the vaccine [33].

In our work, we present the results of creating a candidate mRNA vaccine with an antigen composition recommended by the WHO for use in seasonal vaccines in the northern hemisphere in 2022-2023: A/Wisconsin/588/2019 (H1), A/Darwin/6/2021 (H3), and B/Austria/1359417/2021 (IBV, Victoria lineage) [21]. We used a quadrivalent inactivated split vaccine as a control, which also has the recommended antigen composition. In the main series of experiments, we used vaccines at 1/10 of the human dose (set in the case of split vaccine and assumed for the mRNA vaccine) to allow comparison of platforms when assessing immunogenicity in mice. Both vaccines showed detected immunogenicity using the HAI method. This method is the gold standard for assessing the immunogenicity of influenza vaccines and the efficacy of immunity against novel virus variants [23].

The level of humoral response in HAI to the mRNA vaccine injection was on average 10- 100 times higher than that to the split inactivated vaccine for all subtypes. This correlates with previously obtained data for the combined mRNA vaccine [34]. When immunogenicity was analyzed by ELISA method (against H1N1pdm) for the group of mice immunized with the combined mRNA vaccine, as well as in our case, 10-100 times higher levels of antibodies (depending on the dose of mRNA) were observed compared to this value in the group of animals vaccinated with quadrivalent inactivated vaccine (at a dose of 1.5 µg). The levels of antibody response for the inactivated vaccine obtained in our experiments were in good agreement with the data of the cell-based split vaccine study [35]. For a 1.5 μg dose of each antigen separately, HAI titers comparable to those obtained in our work for the quadrivalent vaccine were obtained (HAI titers to H1N1 averaged about 1:90, to H3N2 averaged about 1:500, and to B (Victoria lineage) averaged about 1:30). One could speculate that this difference between the immune response to the mRNA and the split vaccine could be due to the dose of antigen delivered. Thus, in the case of the inactivated vaccine, the dose of individual antigens was 1,5 μg, whereas for the mRNA composite, the dose of each antigen was 5 μg. Unfortunately, there is no adequate method to quantitatively compare antigen delivered using different vaccine platforms. Delivery of the antigenic protein and its analogue in mRNA form initially yields different pharmacokinetics. According to one of the studies, based on the results of an *in vitro* experiment, 1 pmol of the spike protein was obtained using 6 pmol of mRNA vaccine for 24 hours during transfection of BHK cell culture [36]. According to our earlier experiments, when we delivered 10 μg of mRNA (approximately 20 pmol) encoding the B11 antibody against botulinum toxin we achieved a maximum antibody concentration in mouse serum of 99 ng/mL, which translates to about 2.35 pmol of antibody per average serum volume of an 18-g mouse [22]. Both examples indicate that the amount of antigen produced with the target mRNA is 5-10 times less compared to the amount of mRNA used. Thus, it is more likely that the advantage of mRNA in the context of the generated humoral response is not due to the obviously higher concentration of antigen achieved by the selected doses, but is due to a complex of differences including the mRNA platform itself compared to the delivery of purified antigens used in split vaccines.

As a result of the protective efficacy study using 25 LD_50_ of H1N1 virus (A/Victoria/2570/2019, mouse adapted), we demonstrated the protective effect of both mRNA and split vaccine compositions. In the case of mRNA, no animal deaths were registered during the observation period, whereas in the group of mice that received split vaccine, 3 out of 10 animals died. These data correlated with the dynamics of weight of infected animals (for split vaccine, weight loss was observed from days 3 to 9 of observation), as well as viral load in the lungs - on day 3, the difference in viral load between the groups that received mRNA composition and split vaccine was 10^6^. We were unable to find studies that examined the protectiveness of mRNA and split vaccines in a single animal experiment, but there are studies of split vaccine candidates produced by virus production in the MDCK cell line [35]. On day 4 after 10^7^ CCID_50_ infection with H1N1 influenza virus, a weight loss of up to 10% was observed in the group of mice vaccinated with 5 μg split mono-vaccine, and the amount of virus in the lungs of infected mice isolated on day 6 was 10^2^ virus copies/mL, compared with 10^5^ virus copies/mL in the placebo group. This is indicative that both in our case and in the Zhang et al. study, influenza virus was detected by qPCR in the lungs of animals vaccinated with inactivated split vaccines after infection. In our study, in the mRNA group, the viral load in the lungs was below the detection limit, which is consistent with the data obtained by Arevalo et al. for both 20-valent and monovalent H1-mRNA influenza vaccines [16]. In the lungs of control mice immunized with mRNA vector with luciferase (placebo), the viral load determined by median tissue culture infectious dose (TCID_50_) assay was approximately 5*10^4^ TCID_50_/mL, and in groups of mice immunized with mono H1-mRNA preparation or 20-valent mRNA, the viral load was below the detection limit. As for the results of animal death in the split vaccine group (n = 3), this effect was observed by researchers of adjuvants for influenza vaccines [37]. When intranasally administering a split vaccine, they observed 100% mortality of animals by day 6 after infection with 10 LD_50_ adapted to mice influenza virus A H1N1 A/California/04/2009. On the 3rd day after infection, the virus titer in the bronchoalveolar lavage of infected mice was 6,7 ± 0.1 Log 10 PFU/mL. In general, the death of mice immunized with split influenza vaccines after infection with different doses of influenza viruses is not a new phenomenon. It has been noted in other studies as well [37,38]. Our results for the split-vaccine in terms of the efficacy of protecting vaccinated mice from death are consistent with the published data. However, for preclinical studies of influenza vaccine efficacy, ferrets are the preferred animal model [39].

As a result of using heterologous to the vaccine influenza virus antigens in the HAI test, we demonstrated a broader serum haemagglutinating activity in mice receiving the mRNA composition compared to those receiving the split inactivated influenza vaccine. The immune response to the B/Washington/02/219 strain, which was heterologous to the vaccine strain, was at a level exceeding the lower limit of detection and increased after repeated vaccination. However, in the group of animals immunized with the split vaccine, serum HAI titers were below the LOD at all time points, and no enhancement of immunity by repeated vaccination was observed. The findings indicate a higher potential for the formation of a cross-specific humoral immune response by mRNA compositions. An extended study with four different H1N1 antigens confirmed the findings. Statistically significant differences in serum activity against evolutionarily more distant virus variants (A/California/07/2009pmd, A/Guangdong-Maonan/SWL1536/2019), especially on day 14 after immunization, were detected. However, on day 39, the sera showed high activity against all variants; differences in the level of immune response in hetero- and homologous HAI titers were statistically insignificant, and average serum titers ranged from 1:160 (in the case of the most distant A/California/07/2009pdm) to 1:1 280 (in the case of the epidemic strain A/Moscow/52/22, which circulated in the epidemic season 2022–23 after the vaccine was released). These data correlate with the results of previously published works, which demonstrated the presence of binding and neutralizing antibodies against the H1N1 strain (A/Michigan/45/2015, distant to the vaccine) in the monovalent mRNA vaccine. However, a two-fold reduction in the binding level and the absence of neutralizing antibodies were observed against the second, more distant H1N1 strain, A/Puerto- Rico/08/1934 [16]. We have demonstrated the cross-immunity due to mRNA vaccination against a strain that emerged after updating the vaccine composition for the 2022-2023 season, acquiring 9 substitutions in the amino acid sequence of HA (relative to A/WSN/588/2019). It is important to note that cross-reactivity of the immune response to mRNA vaccine is achieved by the presence of HA mRNA from one strain of influenza subtype in the vaccine.

In summary, the results presented in this study highlight the potential of mRNA-based platforms for the development of influenza vaccines and suggest the need for further research in this area. To fully understand the universality, breadth, and effectiveness of the immune response elicited by mRNA vaccines, it is crucial to conduct additional studies evaluating the immunogenicity and protective efficacy against more diverse strains of H1, H3, and BV *in vivo*. It is also important to validate the data obtained from hemagglutination tests by conducting sera microneutralization assays. Additionally, including a group of mice immunized with non-lethal doses of live virus as a positive control in immunogenicity studies can provide valuable insights into the models of the natural immunity. These findings hold significant implications for the development of mRNA vaccines that offer broader protection against a wide range of influenza strains.

## METHODS

### mRNA production

The pJAZZ-OK-based linear bacterial plasmids (Lucigen) with coding regions of every HAs were used as templates for mRNAs production. DNA cloning procedures were performed as described earlier [22]. The identity of the coding sequences was confirmed by Sanger sequencing. The pDNA for IVT were isolated from the *E.coli* BigEasy™-TSA™ Electrocompetent Cells (Lucigen) using the Plasmid Maxi Kit (QIAGEN). The pDNA was digested using BsmBI-v2 restriction endonuclease (NEB), followed by purification of the product by phenol-chloroform extraction and ethanol precipitation. IVT was performed as described earlier [22]. Briefly, 100-μl reaction volume contained 3 μg of DNA template, 3 μl T7 RNA polymerase (Biolabmix) and 10xBuffer (TriLink), 4 mM trinucleotide cap 1 analog (3′- OMe-m7G)-5′-ppp-5′-(2′-OMeA)pG (Biolabmix), 5 mM m1ΨTP (Biolabmix) replacing UTP, and 5 mM GTP, ATP and CTP. After 2 h incubation at 37 °C, 6 μl DNase I (Thermo Fisher Scientifiс) was added for additional 15 min, followed by mRNA precipitation with 2M LiCl (incubation for 1 h in ice and centrifugation for 30 min at 14,000 g, 4 °C) and carefully washed with 80% ethanol. RNA integrity was assessed by electrophoresis in 8% denaturing PAGE.

### mRNA-LNP assembly

LNP assembly was performed as described earlier [22] with some modifications. In brief, all lipid components were dissolved in ethanol at molar ratios 46.3:9:42.7:1.6 (ionizable lipid:DSPC:cholesterol:PEG-lipid). Acuitas ionizable lipid (ALC- 0315) and PEG-lipid (ALC-0159) were purchased in Cayman Chemicals. The lipid mixture was combined with an acidification buffer of 10 mM sodium citrate (pH 3.0) containing mRNA (0.2 mg/ml) at a volume ratio of 3:1 (aqueous: ethanol) using the NanoAssemblr Ignite device (Precision NanoSystems). The ratio of ionizable nitrogen atoms in the ionizable lipid to the number of phosphate groups in the mRNA (N:P ratio) was set to 6 for each formulation. Formulations were dialyzed against PBS (pH 7.2) in Slide-A-Lyzer dialysis cassettes (Thermo Fisher Scientific) for at least 24 h. Formulations were passed through a 0.22-μm filter and stored at 4 °C (PBS) until use. The diameter and size distribution, zeta potential of the mRNA-LNP were measured using a Zetasizer Nano ZS instrument (Malvern Panalytical) according to user manual.

The mRNA encapsulation efficiency and concentration were determined by SYBR Green dye (SYBR Green I, Lumiprobe) followed by fluorescence measurement. Briefly, mRNA-LNP samples were diluted with TE buffer (pH 8.0) in the absence or presence of 2% Triton-X-100 in a black 96-well plate. Standard mRNA (4 ng/μL) was serially diluted with TE buffer in the absence or presence of 2% Triton-X-100 to generate standard curves. Then the plate was incubated 10 min at room temperature on a rotating shaker (260 rpm) followed by addition of SYBR Green dye (100 times diluted in TE buffer) to each well to bind RNA. Fluorescence was measured at 454 nm excitation and 524 nm emission using Varioscan LUX (Thermo Fisher Scientifiс). The concentrations of mRNA after LNP disruption by Triton-X-100 (C total mRNA) and before LNP disruption (C outside mRNA) were determined using the corresponding standard curves. The concentration of mRNA loaded into the LNP was determined as the difference between the two concentrations multiplied by the dilution factor of the original sample. Encapsulation efficiency was calculated by the formula: (E%)L=L[(C total mRNA) – (C outside mRNA)]/(C total mRNA) ×L100%.

### Cell culture

HEK293 cells (ATCC CRL-1573) were cultured in Dulbecco’s modified Eagle’s medium (DMEM, Paneco) supplemented with glutamine (Gibco), 10% fetal bovine serum (HyClone), 50 U/ml penicillin and 50 μg/ml streptomycin (both from Paneco). Transfection of HEK293 cells with H1N1 HA-encoding mRNA was performed as described previously [22] with minor modification. Briefly, HEK293 cells were plated on a 12-well plate in a density of 2×10^5^cells per well and maintained at 37 °C in 5% CO_2_. The next day the medium was replaced with a fresh DMEM without antibiotics and cells were transfected by HA mRNA using Lipofectamine 3000 reagent (Invitrogen) and Opti-Mem I Reduced Serum Medium (Gibco) in accordance with the manufacturer’s instructions. 24 h after transfection, cells were analyzed by immunocytochemical staining.

### Immunocytochemistry (ICC)

For the analysis of the HA expression in the transfected HEK293 cells, they were fixed in 4% PFA (paraformaldehyde) for 30 min at 40 °C, washed with PBS and permeabilized in 0.1% Triton X-100. The cells were then incubated with primary goat Anti-Influenza A Antibody (Chemicon®, Sigma-Aldrich, #AB1074) in PBS with 0.1%/0.02% BSA/Triton X-100 at 40 °C overnight. The next day, cells were incubated with Donkey Anti-Goat IgG H&L (Alexa Fluor® 488) (ab150133; Abcam) secondary antibodies for 1 h at room temperature. Nuclei were stained with 4,6-diamidino-2-phenylindole (DAPI) (300 nM). Images were acquired on ZOE Fluorescent Cell Imager (Bio-Rad).

### Viruses

Influenza virus (strain H1N1 A/Victoria/2570/2019) propagation in embryonated chicken eggs was performed according to conventional technique [41]. Briefly, specific pathogen-free (SPF) fertilized 9-10 days old chicken eggs were purchased from Nursery “Podmoklovo” (Russia). Presence of the embryo was monitored using an egg candler. Virus inoculation is carried out by injection of virus stock into the allantoic cavity using a needle. After 2 days of incubation at 34 °C, the eggs are cooled for at least 4 h at 4 °C. The eggshell above the air sac and the chorioallantoic membrane are then carefully opened, and the allantoic fluid containing the virus is harvested. The fluid is cleared from debris by centrifugation, aliquoted and transferred to -80 °C for long-term storage. Virus titer was determined by endpoint dilution assay on MDCK cells as previously described [40].

### Animal studies

Females BALB/c mice of 4-5 weeks old were used for the immunogenicity study of HA-mRNA-LNPs and for the viral challenge experiments. Animals were purchased from Stolbovaya breeding and nursery laboratory (Research Center for Biomedical Technologies of FMBA; Russia). All animal experiments were performed in accordance with the Directive 2010/63/EU (Directive 2010/63/EU of the European Parliament and of the Council of 22 September 2010 on the Protection of Animals Used for Scientific Purposes. Off J Eur Communities L276:33–79), FELASA recommendations [41], and the Inter-State Standard of “GLP” [42] approved by the Biomedical Ethics Committee of the Federal Research Centre of Epidemiology and Microbiology named after Honorary Academician N.F. Gamaleya and were performed under Protocol #41 from 6 April 2023. All persons using or caring for animals in research underwent training annually as required by the Biomedical Ethics Committee.

### Immunizations

For mouse intramuscular immunization 100 μl of vaccine was injected in either the left or right hindlimb muscles. Mice received two doses of HA-mRNA-LNPs (altogether or apart) with a 14- or 21-days interval, while the placebo group received PBS. Mice from the positive control group were injected by equal volume (100 μl) of 1/10 human dose of split inactivated influenza vaccine.

### Hemagglutinin inhibition (HAI) assay

Immunogenicity in animal experiments was estimated by hemagglutinin inhibition assay, according to the World Health Organization (WHO)-based HAI protocol [43]. Shortly, mouse sera were treated by a receptor destroying enzyme (RDE produced from Vibrio cholerae was purchased from Denka Seiken Co., Ltd., Tokyo, Japan), then twofold dilutions of treated sera to be tested are made in 96 well plates. The viral antigen was added, and the plate was incubated for 30 minutes at room temperature. Human red blood cells (RBC) type O are then added and the plate incubated for a further 60 minutes at room temperature. If there were antibodies in the serum sample that cross-reacted with the virus, the antibodies would bind to the virus and prevent the virus from haemagglutinating the RBC. After incubation, the HAI titer was red as the highest dilution of serum that inhibited hemagglutination. Antigens of influenza virus for HAI test (A/Darwin/9/2021, A/Victoria/2570/2019, A/Wisconsin/588/2019, B/Austria/1359417/2021) were purchased in LLC "Company for the production of diagnostic drugs" (St-Petersburg, Russia [44]) or propagated in embryonated chicken eggs (B/Washington/02/2019).

### Viral challenge

The lethal infection caused by influenza virus was performed on 4-5-week-old female BALB/c mice. Mouse adapted influenza virus H1N1 A/Victoria/2570/2019 was obtained from the laboratory of molecular biotechnology of the Federal Research Centre of Epidemiology and Microbiology named after Honorary Academician N.F. Gamaleya. Mice were infected intranasally with 50 µL of virus suspension under Zoletil-Xylazine anesthesia. Animals were monitored for clinical symptoms (weight loss, survival) every day through 10 days after the challenge. Assessment of clinical symptoms was carried out in accordance with the scale: score of 0 (no symptoms), score of 1 (mild symptoms), score of 2 (moderate symptoms), score of 3 (severe symptoms = humane endpoint). Time of death was defined as the time at which a mouse was found dead or was euthanized via carbon dioxide asphyxiation followed by cervical dislocation at the endpoint.

### Quantification of virus in infected lungs from mice

Lungs were harvested from mice 3 days post-infection. Following harvest, lungs were weighed, and then homogenized in sterile DMEM with gentamycin to generate a 20% lung-in-medium solution. Total RNA was extracted from lung homogenates using the ExtractRNA Reagent (Eurogen, Moscow, Russia) following the manufacturer’s instructions. Amplification and quantification of influenza A virus RNA were carried out by using a one-step RT-qPCR technique. To perform one-step RT-qPCR was used the reaction mixture containing (for one reaction) 5 pmol of each primer, 3 pmol of probe, 12.5 μl of 2x BioMaster RT-PCR-RT (Biolabmix, Moscow, Russia) and 10 μl of RNA (0,5 μg). The total volume of the one reaction mixture was 25 μl. The primers and probes were designed to target the gene coding M (matrix protein) of influenza A virus, the oligonucleotides were as follows: forward primer – 5’- ATG GAG TGG CTA AAG ACA AGA C -3’, reverse primer - 5’- GCA TTT TGG ACA AAG CGT CTA -3’, probe 5’-FAM - TCC TCG CTC ACT GGG CAC GGT -BHQ1-3’. Amplification was performed using a Real-time CFX96 Touch instrument (Bio-Rad, USA). The conditions of the one-step RT-qPCR reaction were as follows: 50°C for 15 min, 95°C for 5 min, followed by 45 cycles of 95 °C for 10 s and 55 °C for 1 min. The number of copies of viral RNA was calculated using a standard curve generated by amplification of a plasmid cloned DNA template containing the amplified fragment.

## Acknowledgments

We are grateful to Timofey A. Remizov and Anastasia A. Zakharova for their help with the purchase of reagents and paperwork for ongoing projects.

## SUPPLEMENTARY

### Cross-reactivity of immune response in mice after vaccination with 2.5 µg dose of H1 mRNA mono-vaccine

Assuming that over the past four seasons, the most frequent strain changes have occurred in the H1 component of vaccines, we decided to study the cross-reactivity of the immune response against four H1N1 strains after immunization of mice with H1 mRNA (encoding HA A/Wisconsin/588/2019). For this purpose, females of BALB/c mice (n=5 per group) aged 6-7 weeks were immunized twice with an H1-mRNA-LNP (2,5 μg per mouse) with an interval between immunizations of 21 days. The vaccine dose was lower than in experiment with trivalent mRNA vaccine and the time interval between doses was longer by 1 week. The level of immune response was determined by HAI test in the sera of vaccinated mice selected on days 14 and 39 from first vaccination (14 days after prime dose and 18 days after second dose, respectively, Figure 5).

When analyzing immunogenicity in HAI test, antigens of influenza A H1N1 virus strains (A/Wisconsin/588/2019 (clade 6B.1A.5a.2), A/California/07/2009 pdm (clade 6B.1.), A/Guangdong-Maonan/SWL1536/2019 (clade 6B.1A.5a.1), A/Moscow/52/2022 (clade 6B.1A.5a.2a) were used, which belong to different genetic clades of pandemic influenza A H1N1 and have different degrees of homology with the H1 component of the mRNA-IV vaccine by amino acid sequence of HA. Pairwise comparisons of HAI titers to homologous and heterologous antigens revealed a statistically significant decrease in HAI titers to A/California/07/2009 pdm virus antigen in sera on days 14 and 39 after the start of the experiment.

The decrease in the mean titer value for A/California/07/2009 pdm on day 39 was 5,2- fold. A less than 4-fold decrease in mean HAI titer (2,2-fold) was observed to A/Guangdong- Maonan/SWL1536/2019 virus antigen on day 39 compared to the homologous (A/Wisconsin/588/2019), but statistical comparison showed no significant differences (Figure S2). And the mean titer in the serum of mice on day 39 in reaction with the antigen of epidemic strain A/Moscow/52/2022 exceeded that in HAI test with homologous antigen by 0,3 times and the titers to these antigens are not statistically different at both time points (days 14 and 39).

---

**Figure S1.**
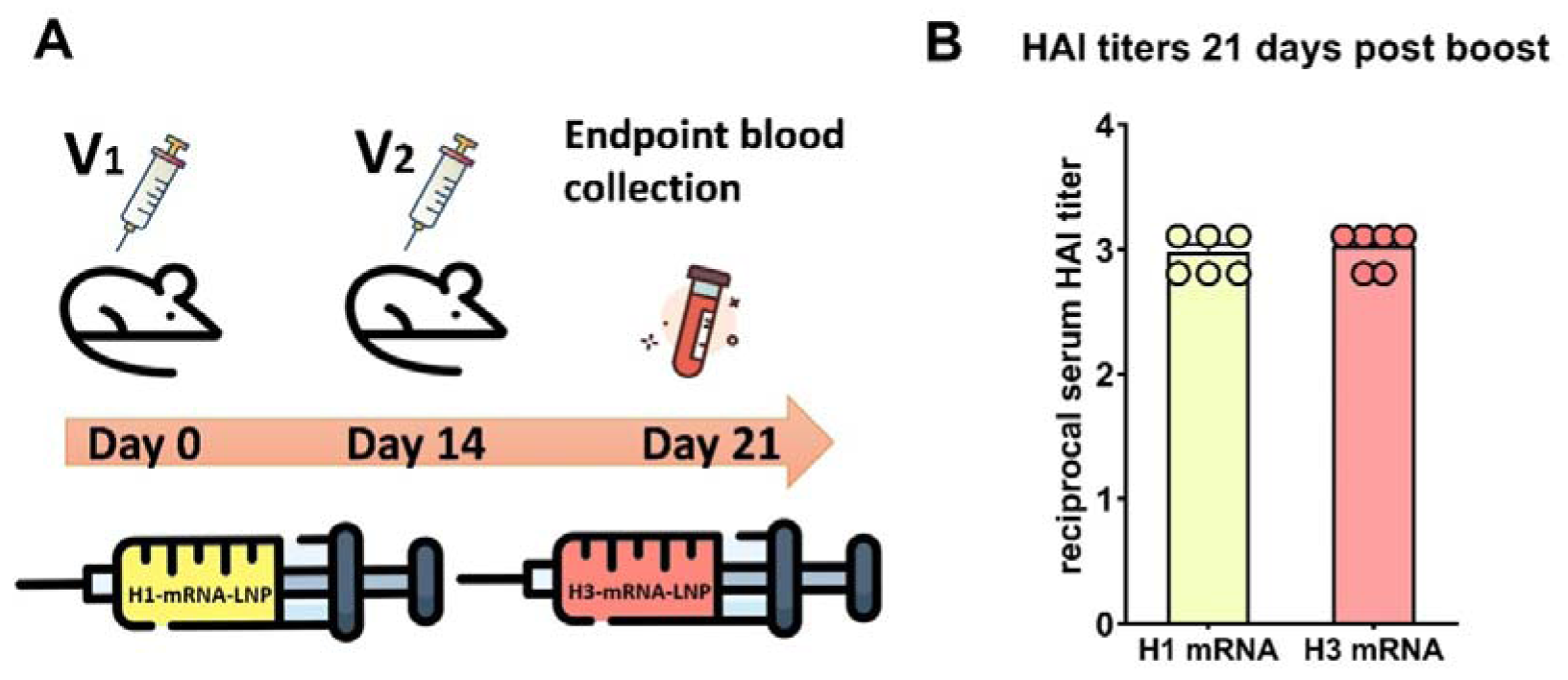
Immunogenicity of H1 and H3 mRNAs mono-vaccines 21 days post second dose. Females of BALB/c mice (n=6 in group) were immunized with H1- or H3 mRNA (10 µg per mice) separately. Second doses were administrated 14 days after the primary dose. HAI titers were determined 7 days post second immunization.

**Figure S2.**
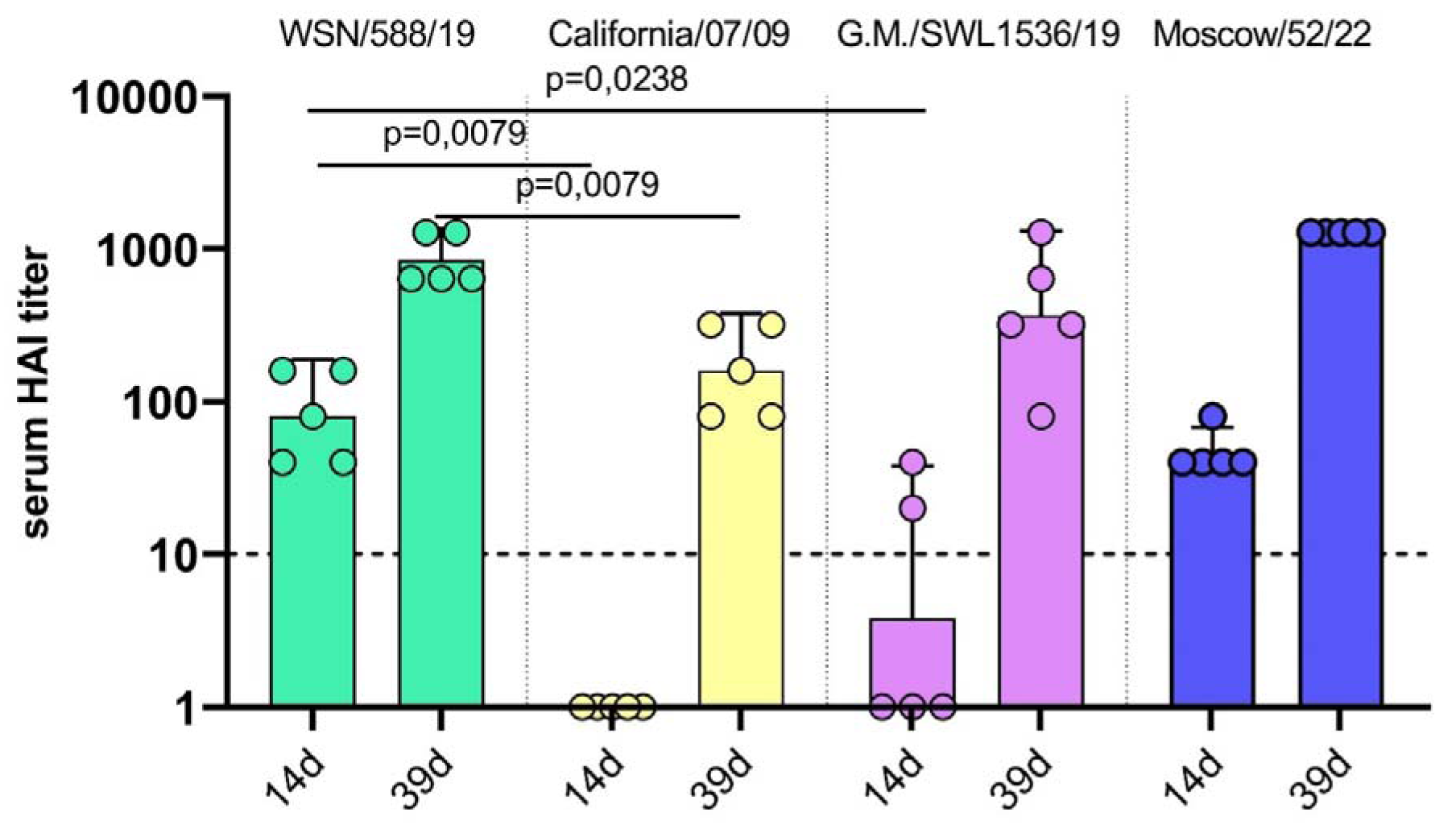
Cross-specific immunogenicity of 2.5 µg H1-mRNA in mice. HAI titers were determined in serum of mice vaccinated with H1-mRNA (n=5 per group) using flu antigens from four different strains of H1N1 influenza A virus (A/Wisconsin/588/2019 (100% amino acid identity of HA sequences), A/California/07/2009 pdm (95.2% HA identity), A/Guangdong- Maonan/SWL1536/2019 (98.6% HA identity), A/Moscow/52/2022 (98.4% HA identity) at 14 and 39 days after first dose. Data are representative of one experiment and shown as geometric means ± SD. Data were compared using a Mann Whitney test.

**Table S1.**
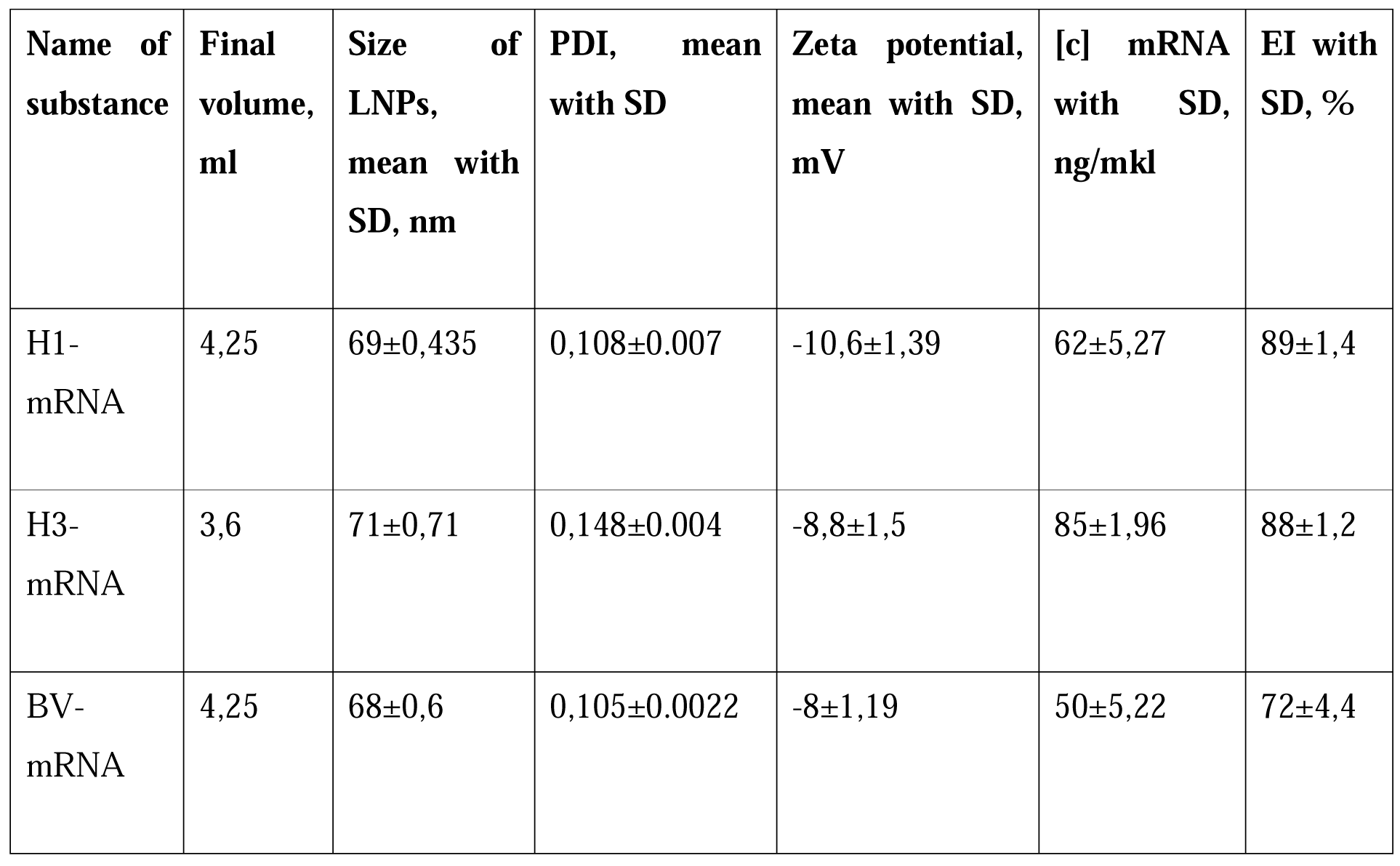
Characteristics of mRNA-LNPs after formulation.

## REFERENCES

1. Krammer, F.; Smith, G.J.D.; Fouchier, R.A.M.; Peiris, M.; Kedzierska, K.; Doherty, P.C.; Palese, P.; Shaw, M.L.; Treanor, J.; Webster, R.G.;, et al. Influenza. Nat. Rev. Dis. Primer 2018, 4, 3, doi:10.1038/s41572-018-0002-y.

2. Factsheet about Seasonal Influenza Available online: https://www.ecdc.europa.eu/en/seasonal-influenza/facts/factsheet (accessed on 27 December 2023).

3. 3. CDC People at High Risk of Flu Available online: https://www.cdc.gov/flu/highrisk/index.htm (accessed on 27 December 2023).

4. Troeger, C.E.; Blacker, B.F.; Khalil, I.A.; Zimsen, S.R.M.; Albertson, S.B.; Abate, D.; Abdela, J.; Adhikari, T.B.; Aghayan, S.A.; Agrawal, S.;, et al. Mortality, Morbidity, and Hospitalisations Due to Influenza Lower Respiratory Tract Infections, 2017: An Analysis for the Global Burden of Disease Study 2017. *Lancet Respir. Med.* 2019, *7*, 69–89, doi:10.1016/S2213-2600(18)30496-X.

5. Sekiya, T.; Ohno, M.; Nomura, N.; Handabile, C.; Shingai, M.; Jackson, D.C.; Brown, L.E.; Kida, H. Selecting and Using the Appropriate Influenza Vaccine for Each Individual. Viruses 2021, 13, 971, doi:10.3390/v13060971.

6. Tenforde, M.W.; Kondor, R.J.G.; Chung, J.R.; Zimmerman, R.K.; Nowalk, M.P.; Jackson, M.L.; Jackson, L.A.; Monto, A.S.; Martin, E.T.; Belongia, E.A.;, et al. Effect of Antigenic Drift on Influenza Vaccine Effectiveness in the United States—2019–2020. Clin. Infect. Dis. 2021, 73, e4244–e4250, doi:10.1093/cid/ciaa1884.

7. Tanner, A.R.; Dorey, R.B.; Brendish, N.J.; Clark, T.W. Influenza Vaccination: Protecting the Most Vulnerable. Eur. Respir. Rev. 2021, 30, 200258, doi:10.1183/16000617.0258-2020.

8. CDC Key Facts About Influenza (Flu) Available online: https://www.cdc.gov/flu/about/keyfacts.htm (accessed on 27 December 2023).

9. Ferdinands, J.M.; Thompson, M.G.; Blanton, L.; Spencer, S.; Grant, L.; Fry, A.M. Does Influenza Vaccination Attenuate the Severity of Breakthrough Infections? A Narrative Review and Recommendations for Further Research. Vaccine 2021, 39, 3678–3695, doi:10.1016/j.vaccine.2021.05.011.

10. Ferdinands, J.M.; Olsho, L.E.W.; Agan, A.A.; Bhat, N.; Sullivan, R.M.; Hall, M.; Mourani, P.M.; Thompson, M.; Randolph, A.G. Effectiveness of Influenza Vaccine Against Life-Threatening RT-PCR-Confirmed Influenza Illness in US Children, 2010–2012. *J. Infect. Dis.* 2014, *210*, 674–683, doi:10.1093/infdis/jiu185.

11. Rondy, M.; El Omeiri, N.; Thompson, M.G.; Levêque, A.; Moren, A.; Sullivan, S.G. Effectiveness of Influenza Vaccines in Preventing Severe Influenza Illness among Adults: A Systematic Review and Meta-Analysis of Test-Negative Design Case-Control Studies. J. Infect. 2017, 75, 381–394, doi:10.1016/j.jinf.2017.09.010.

12. Xie, H.; Wan, X.-F.; Ye, Z.; Plant, E.P.; Zhao, Y.; Xu, Y.; Li, X.; Finch, C.; Zhao, N.;Kawano, T.;, et al. H3N2 Mismatch of 2014–15 Northern Hemisphere Influenza Vaccines and Head-to-Head Comparison between Human and Ferret Antisera Derived Antigenic Maps. Sci. Rep. 2015, 5, 15279, doi:10.1038/srep15279.

13. Erbelding, E.J.; Post, D.J.; Stemmy, E.J.; Roberts, P.C.; Augustine, A.D.; Ferguson, S.; Paules, C.I.; Graham, B.S.; Fauci, A.S. A Universal Influenza Vaccine: The Strategic Plan for the National Institute of Allergy and Infectious Diseases. J. Infect. Dis. 2018, 218, 347– 354, doi:10.1093/infdis/jiy103.

14. Polack, F.P.; Thomas, S.J.; Kitchin, N.; Absalon, J.; Gurtman, A.; Lockhart, S.; Perez, J.L.; Pérez Marc, G.; Moreira, E.D.; Zerbini, C.;, et al. Safety and Efficacy of the BNT162b2 mRNA Covid-19 Vaccine. N. Engl. J. Med. 2020, 383, 2603–2615, doi:10.1056/NEJMoa2034577.

15. Baden, L.R.; El Sahly, H.M.; Essink, B.; Kotloff, K.; Frey, S.; Novak, R.; Diemert, D.; Spector, S.A.; Rouphael, N.; Creech, C.B.;, et al. Efficacy and Safety of the mRNA-1273 SARS-CoV-2 Vaccine. N. Engl. J. Med. 2021, 384, 403–416, doi:10.1056/NEJMoa2035389.

16. Arevalo, C.P.; Bolton, M.J.; Le Sage, V.; Ye, N.; Furey, C.; Muramatsu, H.; Alameh, M.-G.; Pardi, N.; Drapeau, E.M.; Parkhouse, K.;, et al. A Multivalent Nucleoside-Modified mRNA Vaccine against All Known Influenza Virus Subtypes. Science 2022, 378, 899–904, doi:10.1126/science.abm0271.

17. Nachbagauer, R.; Liu, W.-C.; Choi, A.; Wohlbold, T.J.; Atlas, T.; Rajendran, M.; Solórzano, A.; Berlanda-Scorza, F.; García-Sastre, A.; Palese, P.;, et al. A Universal Influenza Virus Vaccine Candidate Confers Protection against Pandemic H1N1 Infection in Preclinical Ferret Studies. Npj Vaccines 2017, 2, 26, doi:10.1038/s41541-017-0026-4.

18. Pardi, N.; Parkhouse, K.; Kirkpatrick, E.; McMahon, M.; Zost, S.J.; Mui, B.L.; Tam, Y.K.; Karikó, K.; Barbosa, C.J.; Madden, T.D.;, et al. Nucleoside-Modified mRNA Immunization Elicits Influenza Virus Hemagglutinin Stalk-Specific Antibodies. Nat. Commun. 2018, 9, 3361, doi:10.1038/s41467-018-05482-0.

19. Bahl, K.; Senn, J.J.; Yuzhakov, O.; Bulychev, A.; Brito, L.A.; Hassett, K.J.; Laska, M.E.; Smith, M.; Almarsson, Ö.; Thompson, J.;, et al. Preclinical and Clinical Demonstration of Immunogenicity by mRNA Vaccines against H10N8 and H7N9 Influenza Viruses. Mol. Ther. 2017, 25, 1316–1327, doi:10.1016/j.ymthe.2017.03.035.

20. Feldman, R.A.; Fuhr, R.; Smolenov, I.; (Mick) Ribeiro, A.; Panther, L.; Watson, M.; Senn, J.J.; Smith, M.; Almarsson, Lrn; Pujar, H.S.;, et al. mRNA Vaccines against H10N8 and H7N9 Influenza Viruses of Pandemic Potential Are Immunogenic and Well Tolerated in Healthy Adults in Phase 1 Randomized Clinical Trials. Vaccine 2019, 37, 3326–3334, doi:10.1016/j.vaccine.2019.04.074.

21. Recommendations Announced for Influenza Vaccine Composition for the 2022-2023 Northern Hemisphere Influenza Season Available online: https://www.who.int/news/item/25-02-2022-recommendations-announced-for-influenza-vaccine-composition-for-the-2022-2023-northern-hemisphere-influenza-season (accessed on 27 December 2023).

22. Panova, E.A.; Kleymenov, D.A.; Shcheblyakov, D.V.; Bykonia, E.N.; Mazunina, E.P.; Dzharullaeva, A.S.; Zolotar, A.N.; Derkaev, A.A.; Esmagambetov, I.B.; Sorokin, I.I.;, et al. Single-Domain Antibody Delivery Using an mRNA Platform Protects against Lethal Doses of Botulinum Neurotoxin A. Front. Immunol. 2023, 14, 1098302, doi:10.3389/fimmu.2023.1098302.

23. L’vov, D.K.; Burtseva, E.I.; Kolobukhina, L.V.; Fedyakina, I.T.; Bovin, N.V.; Ignatjeva, A.V.; Krasnoslobodtsev, K.G.; Feodoritova, E.L.; Trushakova, S.V.; Breslav, N.V.;, et al. Peculiarities of the Influenza and ARVI Viruses Circulation during Epidemic Season 2019– 2020 in Some Regions of Russia. Probl. Virol. 2021, 65, 335–349, doi:10.36233/0507-4088-2020-65-6-4.

24. Both, G.W.; Sleigh, M.J.; Cox, N.J.; Kendal, A.P. Antigenic Drift in Influenza Virus H3 Hemagglutinin from 1968 to 1980: Multiple Evolutionary Pathways and Sequential Amino Acid Changes at Key Antigenic Sites. J. Virol. 1983, 48, 52–60, doi:10.1128/jvi.48.1.52-60.1983.

25. Hirst, G.K.; Gotlieb, T. THE EXPERIMENTAL PRODUCTION OF COMBINATION FORMS OF VIRUS. J. Exp. Med. 1953, 98, 41–52, doi:10.1084/jem.98.1.41.

26. Gibbs, M.J.; Armstrong, J.S.; Gibbs, A.J. Recombination in the Hemagglutinin Gene of the 1918 “Spanish Flu.” Science 2001, 293, 1842–1845, doi:10.1126/science.1061662.

27. Influenza (Seasonal) Available online: https://www.who.int/news-room/fact-sheets/detail/influenza-(seasonal) (accessed on 28 December 2023).

28. Recommendations for Influenza Vaccine Composition Available online: https://www.who.int/teams/global-influenza-programme/vaccines/who-recommendations (accessed on 27 December 2023).

29. Krammer, F. The Human Antibody Response to Influenza A Virus Infection and Vaccination. Nat. Rev. Immunol. 2019, 19, 383–397, doi:10.1038/s41577-019-0143-6.

30. Ueno, H. Tfh Cell Response in Influenza Vaccines in Humans: What Is Visible and What Is Invisible. Curr. Opin. Immunol. 2019, 59, 9–14, doi:10.1016/j.coi.2019.02.007.

31. Chaudhary, N.; Weissman, D.; Whitehead, K.A. mRNA Vaccines for Infectious Diseases: Principles, Delivery and Clinical Translation. Nat. Rev. Drug Discov. 2021, 20, 817–838, doi:10.1038/s41573-021-00283-5.

32. Wong, S.-S.; Webby, R.J. Traditional and New Influenza Vaccines. Clin. Microbiol. Rev. 2013, 26, 476–492, doi:10.1128/CMR.00097-12.

33. Vogel, A.B.; Kanevsky, I.; Che, Y.; Swanson, K.A.; Muik, A.; Vormehr, M.; Kranz, L.M.; Walzer, K.C.; Hein, S.; Güler, A.;, et al. BNT162b Vaccines Protect Rhesus Macaques from SARS-CoV-2. Nature 2021, 592, 283–289, doi:10.1038/s41586-021-03275-y.

34. Freyn, A.W.; Ramos Da Silva, J.; Rosado, V.C.; Bliss, C.M.; Pine, M.; Mui, B.L.; Tam, Y.K.; Madden, T.D.; De Souza Ferreira, L.C.; Weissman, D.;, et al. A Multi-Targeting, Nucleoside-Modified mRNA Influenza Virus Vaccine Provides Broad Protection in Mice. Mol. Ther. 2020, 28, 1569–1584, doi:10.1016/j.ymthe.2020.04.018.

35. Zhang, J.; Nian, X.; Liu, B.; Zhang, Z.; Zhao, W.; Han, X.; Ma, Y.; Jin, D.; Ma, H.; Zhang, Q.;, et al. Development of MDCK-Based Quadrivalent Split Seasonal Influenza Virus Vaccine with High Safety and Immunoprotection: A Preclinical Study. Antiviral Res. 2023, 216, 105639, doi:10.1016/j.antiviral.2023.105639.

36. Sutton, W.J.H.; Branham, P.J.; Williamson, Y.M.; Cooper, H.C.; Najjar, F.N.; Pierce-Ruiz, C.L.; Barr, J.R.; Williams, T.L. Quantification of SARS-CoV-2 Spike Protein Expression from mRNA Vaccines Using Isotope Dilution Mass Spectrometry. Vaccine 2023, 41, 3872– 3884, doi:10.1016/j.vaccine.2023.04.044.

37. Van De Sandt, C.E.; Kreijtz, J.H.C.M.; Geelhoed-Mieras, M.M.; Trierum, S.E.V.; Nieuwkoop, N.J.; Van De Vijver, D.A.M.C.; Fouchier, R.A.M.; Osterhaus, A.D.M.E.; Morein, B.; Rimmelzwaan, G.F. Novel G3/DT Adjuvant Promotes the Induction of Protective T Cells Responses after Vaccination with a Seasonal Trivalent Inactivated Split-Virion Influenza Vaccine. Vaccine 2014, 32, 5614–5623, doi:10.1016/j.vaccine.2014.08.003.

38. Boravleva, E.Y.; Lunitsin, A.V.; Kaplun, A.P.; Bykova, N.V.; Krasilnikov, I.V.; Gambaryan, A.S. Immune Response and Protective Efficacy of Inactivated and Live Influenza Vaccines Against Homologous and Heterosubtypic Challenge. Biochem. Mosc. 2020, 85, 553–566, doi:10.1134/S0006297920050041.

39. Influenza Vaccines - Non-Clinical and Clinical Module - Scientific Guideline | European Medicines Agency Available online: https://www.ema.europa.eu/en/influenza-vaccines-non-clinical-and-clinical-module-scientific-guideline#ema-inpage-item-topics (accessed on 27 December 2023).

40. Kaufmann, L.; Syedbasha, M.; Vogt, D.; Hollenstein, Y.; Hartmann, J.; Linnik, J.E.; Egli, A. An Optimized Hemagglutination Inhibition (HI) Assay to Quantify Influenza-Specific Antibody Titers. J. Vis. Exp. 2017, 55833, doi:10.3791/55833.

41. FELASA working group on revision of guidelines for health monitoring of rodents and rabbits; Mähler (Convenor), M.; Berard, M.; Feinstein, R.; Gallagher, A.; Illgen-Wilcke, B.; Pritchett-Corning, K.; Raspa, M. FELASA Recommendations for the Health Monitoring of Mouse, Rat, Hamster, Guinea Pig and Rabbit Colonies in Breeding and Experimental Units. *Lab. Anim.* 2014, *48*, 178–192, doi:10.1177/0023677213516312.

42. *GOST 33044-2014;* 2015;

43. Cox, N., Webster, R. and Stohr, K. WHO Manual on Animal Influenza Diagnosis and Surveillance. World Health Organization Department Communicable Disease Surveillance and Response.

44. Главная - ООО «Предприятие По Производству Диагностических Препаратов» (ООО «ППДП») Available online: http://ppdp-spb.com/ (accessed on 28 December 2023).

